# The Xrn1-Rpb4 axis controls mRNA synthesis and imprinting

**DOI:** 10.64898/2026.11.16.766666

**Authors:** Ana I. Garrido-Godino, M. Cintas-Galán, A. Jordán-Pla, JE. Pérez-Ortín, Francisco Navarro

## Abstract

Co-transcriptional binding (imprinting) of mRNAs by the Rpb4 subunit of RNA polymerase II (RNA pol II) coordinates transcription with cytoplasmic mRNA fates. However, the genomic features of Rpb4-target genes and their cross-talk with decay factors remain unclear. In this study, in *Saccharomyces cerevisiae*, we demonstrate that genes producing Rpb4-imprinted mRNAs exhibit higher TR paired with lower levels of RNA pol II backtracking. Notably, these loci act as specific genomic targets for the 5’-3’ exoribonuclease Xrn1, that has also been shown to act as transcriptional activator in a set of yeast genes, displaying prominent Xrn1 enrichment along their gene bodies. In accordance, *XRN1* deletion alters global RNA pol II occupancy, leads to a decrease in TR and reduces growth severely in a *rpb4*Δ background. Furthermore, we show that Rpb4 physically interacts with Xrn1 in a strictly RNA-dependent-imprinting manner. Together, our findings uncover a functional Rpb4–Xrn1 axis that connects efficient transcription elongation with mRNA imprinting and turnover.

## INTRODUCTION

Gene expression in eukaryotes is traditionally viewed as a sequence of compartmentalized steps, starting with transcription in the nucleus and culminating in translation and mRNA decay in the cytoplasm. However, emerging evidences during de last decade demonstrate that these processes are functionally and physically coupled (Begley et al., 2019, 2021; Blasco-Moreno et al., 2019; Bryll & Peterson, 2023; Garrido-Godino et al., 2016, 2021; Goler-Baron et al., 2008; Haimovich et al., 2013; Harel-Sharvit et al., 2010; Hartenian & Glaunsinger, 2019; Kelbert et al., 2024; Medina et al., 2014; Timmers & Tora, 2018). Thereby, total cellular mRNA concentrations are maintained in a tightly regulated dynamic equilibrium, termed mRNA homeostasis, through an intricate mechanism known as mRNA cross-talk or mRNA buffering (García-Martínez et al., 2021; Timmers & Tora, 2018). This integrative cross-talk establishes continuous, bidirectional communication and reciprocal feedback loops between the nuclear transcription machinery and cytoplasmic degradation pathways. This coordination ensures that perturbations in mRNA stability are compensated by adjustments in transcription rates (TR), and *vice versa*, enabling cells to rapidly adapt their transcriptome during metabolic transitions and acute environmental stress conditions (Blasco-Moreno et al., 2019; Chávez et al., 2016; Duncan-Lewis et al., 2021; Pérez-Ortín et al., 2019).

A principal mechanism driving this functional connection in the nucleus to cytoplasm direction is mRNA imprinting, a co-transcriptional process in which nuclear factors bind to the nascent transcript during synthesis and remain attached during its export to the cytoplasm. By remaining associated with the mature mRNA, these imprinted factors directly govern post-transcriptional outcomes, including translational efficiency, subcellular localization, and decay rates (Choder, 2011; Dahan & Choder, 2013; Duek et al., 2018; Garrido-Godino et al., 2016; Goler-Baron et al., 2008; Gupta et al., 2016; Jordán-Pla et al., 2025; Kelbert et al., 2024). Through this mechanism, the events of nuclear transcription directly predetermine the cytoplasmic fate, including lifespan, of the transcript.

Central to this regulatory system is a specialized class of multifunctional factors, frequently referred to as “synthegradases”, which operate directly in both nuclear synthesis and cytoplasmic decay pathways (Haimovich et al., 2013; Pérez-Ortín, Alepuz, et al., 2013). Prominent examples of them in the yeast *S. cerevisiae* include the Rpb4/7 subcomplex of RNA polymerase II (Garrido-Godino et al., 2016, 2021; Goler-Baron et al., 2008; Harel-Sharvit et al., 2010; Lotan et al., 2005), the multifunctional Ccr4-Not complex (Begley et al., 2019; Gupta et al., 2016; Villanyi et al., 2014), and major components of the cytoplasmic decaysome, most notably the 5’→ 3’ exoribonuclease Xrn1 along with decapping factors (Fischer et al., 2020; Haimovich et al., 2013; Medina et al., 2014), all of which bridge nuclear and cytoplasmic processes to regulate gene expression globally.

RNA polymerase II (RNA pol II) serves as the primary engine for eukaryotic transcription, with its non-essential subcomplex Rpb4/7 acting as a key functional bridge between transcription and downstream mRNA fate. Beyond its role in promoter open-complex formation and transcription elongation (Babbarwal et al., 2014; Edwards et al., 1991; Orlicky et al., 2001; Runner et al., 2008; Schulz et al., 2014; Verma-Gaur et al., 2008), Rpb4 is loaded onto nascent mRNAs during transcription in an imprinting process (Duek et al., 2018; Garrido-Godino et al., 2016, 2021; Goler-Baron et al., 2008). Rpb4 remains associated with the transcript during its export to the cytoplasm, where it stimulates mRNA translation and orchestrates subsequent decay by recruiting the deadenylation and decapping machinery (Garrido-Godino et al., 2022). Consequently, Rpb4 —and the Rpb4/7 dimer— act as key elements in mRNA imprinting, functionally coordinating transcription with cytoplasmic fates by interacting with diverse macromolecular complexes. Specifically, Rpb4 physically associates with the Ccr4-Not complex to regulate transcription elongation (Babbarwal et al., 2014) and the Pat1/Lsm1-7 module (Lotan et al., 2005), promoting both Ccr4-targeted mRNA deadenylation and post-deadenylation decay steps like decapping (Bloom et al., 2013; Lotan et al., 2005). Beyond mRNA degradation, Rpb4/7 cooperates with specialized proteins to modulate mRNA export (via Npl3 (Dori-Bachash et al., 2011; Farago et al., 2003)) and translation (via eIF3 components (Harel-Sharvit et al., 2010; Villanyi et al., 2014)). In addition, Rpb4 interacts physically and genetically with the RNA binding protein Puf3 to control de mRNA fate of its mRNA targets (Garrido-Godino et al., 2021). Furthermore, Rpb4 cooperates with Sfp1, a zing-finger factor also participating in the cross-talk between the synthesis and decay of specific mRNAs (Kelbert et al., 2024). Interestingly, several of these factors shuttle between cellular compartments, participate in transcription elongation, or associate with chromatin to contact nascent transcripts co-transcriptionally making it possible the feed-back of cytoplasmic information to nuclear processes (Babbarwal et al., 2014; Garrido-Godino et al., 2021; Miller & Reese, 2012; Villanyi et al., 2014).

In the cytoplasm, the highly conserved 5′-3′ exonuclease Xrn1 serves as the primary driver of bulk mRNA degradation following deadenylation and decapping by the Dcp1/Dcp2 complex (Langeberg et al., 2020; Larimer et al., 1992; Parker & Song, 2004; Stevens, 2001). Operating processively and often co-translationally following the last translating ribosome (Pelechano et al., 2015), Xrn1 also participates in non-coding RNA and hypomodified transfer RNA (tRNA) decay, rRNA processing, and quality control surveillance (Chernyakov et al., 2008; Geerlings et al., 2000; Parker, 2012; Van Dijk et al., 2011). Intriguingly, Xrn1 function is not confined to cytoplasmic clearance; it also feedback-regulates nuclear transcription playing a fundamental role in transcript buffering (Begley et al., 2019, 2021; Blasco-Moreno et al., 2019; Fischer et al., 2020; García-Martínez, José et al., 2021; Haimovich et al., 2013; Medina et al., 2014). Xrn1 undergoes nucleocytoplasmic shuttling regulated by the importin Kap120 via two nuclear localization signals (NLS1 and NLS2). This process is controlled by the degrading mRNA itself, which binds to the enzyme and masks NLS1 to prevent its nuclear import until cytoplasmic decay has concluded (Chattopadhyay et al., 2022).

Once inside the nucleus, Xrn1 binds to promoters (preferably ∼30 bp upstream of the transcription start site) and gene bodies, where it directly stimulates transcription initiation and elongation by increasing RNA pol II processivity and preventing unproductive pausing or backtracking, a transcriptional arrest phenomenon where the enzyme reverses along the DNA template and requires cleavage factor TFIIS for rescue, acting in a parallel and independent manner from the Ccr4-Not complex (Begley et al., 2019, 2021; Fischer et al., 2020; García-Martínez et al., 2021; Haimovich et al., 2013; Medina et al., 2014). Moreover, Xrn1 accumulates at gene 3′ ends and polyadenylation sites (PAS), suggesting a direct involvement in transcription termination and elongation control (Begley et al., 2021; Fischer et al., 2020; García-Martínez et al., 2021). Furthermore, this nuclear transport of Xrn1 is required for efficient mRNA degradation in the cytoplasm through the imprinting of nascent transcripts during their synthesis (Jordán-Pla et al., 2025). In addition to connecting transcription and degradation, Xrn1 activates the translation and endoplasmic reticulum localization of specific transcripts encoding membrane proteins with structured 5’ UTR regions through its interaction with the initiation factor eIF4G (Blasco-Moreno et al., 2019). Consequently, Xrn1 functions as a master coordinator that fully integrates the three main stages of gene expression to enable a rapid reorganization of the transcriptome and proteome in response to environmental changes and stress conditions.

Given that Rpb4 interacts with numerous components of both the mRNA synthesis and degradation machineries, we propose that it may function as a central regulator of mRNA cross-talk. Despite well-documented roles for both Rpb4 and Xrn1 in maintaining mRNA homeostasis, the precise genetic, functional, and physical crosstalk between these two central factors remains incompletely understood.

In this study, we demonstrate a robust functional and physical interplay between Rpb4 and Xrn1 in *S. cerevisiae.* We show that simultaneous loss of *RPB4* and *XRN1* leads to a severe synthetic growth defect, reflecting the importance of genetic buffering between transcription and decay pathways. Notably, the study reveals that genes which mRNAs are imprinted by Rpb4 are genomic targets of Xrn1. Furthermore, using chromatin fractionation and UV-crosslinking, we establish that Rpb4 and Xrn1 bind chromatin and RNA independently. Crucially, co-immunoprecipitation experiments demonstrate that Rpb4 physically contacts Xrn1 imprinting in a strictly RNA-dependent manner, while Rpb4 is also required to maintain Xrn1 protein integrity. Together, these findings uncover an integrated regulatory axis in which Rpb4 and Xrn1 physically and functionally align mRNA synthesis with decay.

## RESULTS

### Genes whose mRNA are imprinted by Rpb4 exhibit higher Rpb3 occupancy compared to other genes

Rpb4 has been shown to imprint mRNA co-transcriptionally to regulate the mRNA fate (Duek et al., 2018; Garrido-Godino et al., 2016, 2022; Goler-Baron et al., 2008; Lotan et al., 2005). Using RIP-Seq experiments, we previously demonstrated that Rpb4 binds to roughly 1,500 mRNAs under permissive conditions (Garrido-Godino et al., 2021). Nevertheless, the genomic features of the genes encoding these Rpb4-bound mRNAs (hereafter referred to as Rpb4-target genes) remain unknown. In order to characterize those genes, we determined RNA pol II occupancy by ChIP-Seq analysis using an antibody against the Rpb3 subunit in a wild-type strain grown at 30°C in YPD medium. Using this approach, we compared the recruitment of RNA pol II (Rpb3-ChIP-Seq signal) at genes whose mRNAs are imprinted by Rpb4 (Rpb4-target genes) versus the rest of the genome. As shown in the metagene plot (Figure 1), Rpb4-target genes displayed increased RNA pol II recruitment than the rest of genes. The shape of the Rpb3 occupancy profile was similar for all type of genes. These findings suggest that Rpb4-target genes exhibit either an average transcriptional elongation but a higher TR, or alternatively, a higher density of non-elongating (backtracked or paused) RNA polymerases.

**Figure 1.**
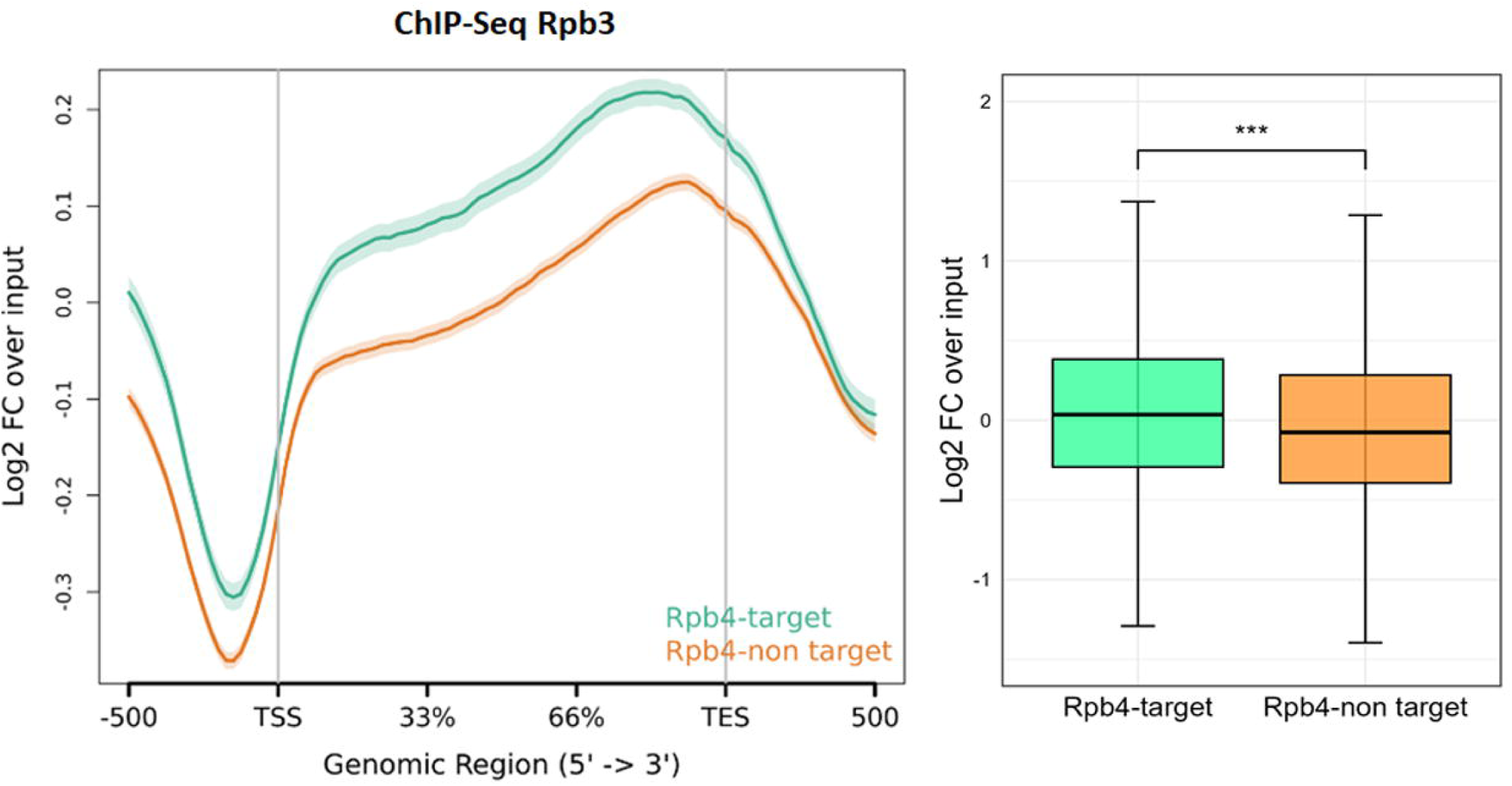
ChIP-Seq analysis of Rpb3 occupancy in a wild-type strain. The plot represents the input-normalized average metagene profile of Rpb3 over gene bodies and flanking regions of all protein-coding genes, categorized into Rpb4-target genes and non-target genes. The profile shows the log_2_ fold change between the Rpb3 ChIP and input signals. Data represent the mean of two biological replicates. TSS, transcription start site; TES, transcription end site. Asterisks indicate statistically significant differences (* P < 0.05, ** P < 0.01, and *** P < 0.001; Wilcoxon rank-sum test). TSS, transcription start site; TES, transcription end site.

### Rpb4-target genes show higher transcription rate and lower backtracking

In line with previous results, we asked whether Rpb4-target genes could have higher TR. To do so, we re-analysed previously published BioGRO-Seq data (Begley et al., 2021). BioGRO-Seq detects only true active elongating RNA pol II, thus reflecting nascent TR (Pérez-Ortín, Medina, et al., 2013). As shown in Figure 2A, genes encoding Rpb4-bound mRNAs displayed higher TR than the rest of the genes. These results are consistent with the higher RNA pol II occupancy determined by ChIP-Seq (Figure 1). Notably, when plotted the metagene of Rpb3 occupancy relative to the BioGRO-Seq signal (Figure 2B), a slightly decreased, but statistically significant, ratio for Rpb4-target genes (green) was observed compared to the rest of the genes (orange) (Figure 2B-Right panel). Given that the ratio Rpb3/BioGRO-Seq represents the enrichment in non-active elongating RNA pol II onto gene bodies, these findings indicate that Rpb4-target genes exhibit lower levels of backtracking than the rest of the genes.

**Figure 2.**
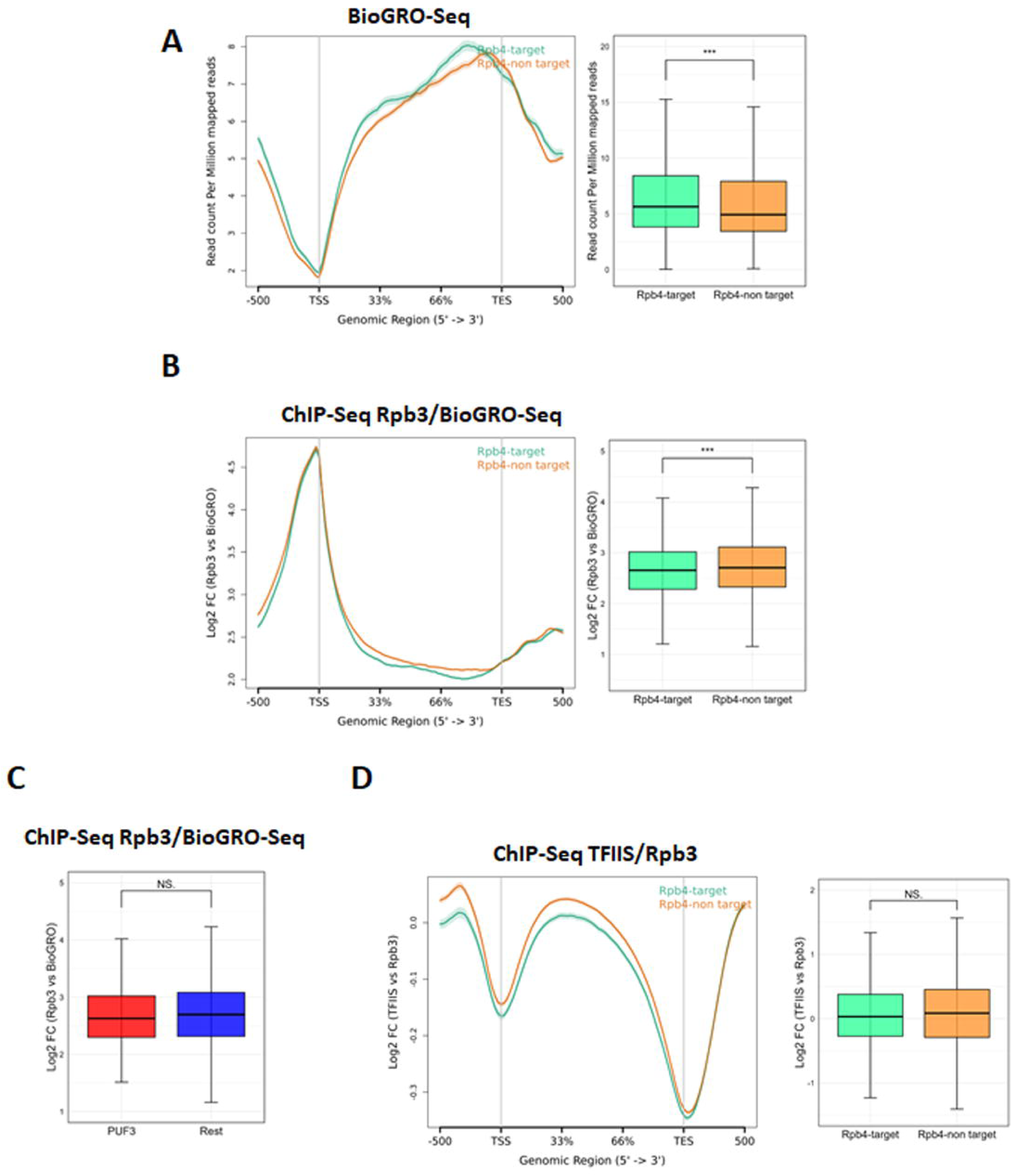
Rpb4-target genes show lower backtracking levels than non-target genes. **(A)** *Left:* Metagene profile of BioGRO-Seq data over gene bodies and flanking regions for protein-coding genes in the wild-type strain, categorized into Rpb4-target and non-target genes. *Right:* Boxplot of the data represented in the left panel. **(B)** *Left:* Average metagene profile showing the log_2_ fold change of Rpb3 occupancy over BioGRO-Seq signal across gene bodies and flanking regions. *Right:* Boxplot of the data represented in the left panel. **(C)** Boxplot showing the Rpb3/BioGRO-Seq signal ratio for Puf3-target genes versus non-target genes. **(D)** Metagene profile of the TFIIS/Rpb3 ratio (log_2_ fold change of TFIIS ChIP-Seq over Rpb3 ChIP-Seq) across Rpb4-target and non-target genes. *Right:* Boxplot of gene body signal from the left panel.Data represent the mean of two biological replicates. Asterisks indicate statistically significant differences (* P < 0.05, ** P < 0.01, and *** P < 0.001; Wilcoxon rank-sum test). TSS, transcription start site; TES, transcription end site.

We have previously demonstrated that a small subset (10%) of these Rpb4-bound mRNAs is also targeted by the RNA-binding protein Puf3 (Garrido-Godino et al., 2021). To determine whether these observations were uniquely correlated to Rpb4, we next evaluated the Rpb3/BioGRO-Seq ratio specifically for the Puf3-target genes (Freeberg et al., 2013; Gerber et al., 2004; Gupta et al., 2014; Kershaw et al., 2015; Wilinski et al., 2017), using the target definitions previously established in our workflow (Garrido-Godino et al., 2021). As shown in Figure 2C, Rpb3/BioGRO-Seq ratio for Puf3-target genes was similar to the rest of the genes, suggesting that the lower Rpb3/BioGRO-Seq ratio shown above, was indeed a specific feature of Rpb4-target genes.

Taken together, these data lead point to Rpb4-target genes constituting a distinct set of genes characterized by high TR accompanied by reduced backtracking. In order to confirm the decrease in backtracking of RNA pol II for Rpb4-target genes, we analysed the occupancy of the Transcription Factor IIS (TFIIS), an RNA cleavage factor that alleviates backtracking by promoting intrinsic RNA cleavage of the backtracked transcript, allowing transcription to resume (Cheung & Cramer, 2011; Gomez-Herreros, de Miguel-Jimenez, Millan-Zambrano, et al., 2012; Gomez-Herreros, de Miguel-Jimenez, Morillo-Huesca, et al., 2012; Izban & Luse, 1992). For that purpose we used a previously published datasheet (Begley et al., 2019). Consistent with the lower Rpb3/BioGRO-Seq ratio, the Rpb4-target gene subset also exhibited a slightly lower, although not statistically significant TFIIS/Rpb3 ratio (Figure 2D).

### Rpb4-target genes are genomic targets of the Xrn1 regulator

Considering that Rpb4/7 drives mRNA imprinting to functionally coordinate transcription with cytoplasmic fates—such as decay, export, and translation—through interactions with diverse macromolecular complexes, we hypothesize that Rpb4 acts as a central hub regulating overall mRNA levels through its interaction with specific RBPs.

To test this hypothesis, we performed ChIP-Seq analyses to profile the recruitment of the decay factors Xrn1, Ccr4, and Dhh1 to Rpb4-target genes, as well as to the rest of genes (Figure 3) using strains expressing TAP-tagged versions of Xrn1, Ccr4 and Dhh1 proteins. As shown in Figure 3A, metagene analysis revealed an increased occupancy of Xrn1 across the entire gene body of Rpb4-target genes with a different profile than Rpb4-non target genes. Consistent with this finding, previous studies have demonstrated the binding of Xrn1 to both promoters and the gene body of highly transcribed genes (García-Martínez et al., 2021; Haimovich et al., 2013; Harlen & Churchman, 2017; Medina et al., 2014). However, the Ccr4 deadenylase (Babbarwal et al., 2014; Begley et al., 2019; Tucker et al., 2002) showed only a slightly higher occupancy at Rpb4-target (Figure 3B). Notably, occupancy of Dhh1 RNA helicase associated with transcription and cytoplasmic decay (Haimovich et al., 2013) decreased at Rpb4-target genes (Figure 3C). Taken together, these findings reveal a specific recruitment pattern of Rpb4 target genes that supports our hypothesis of Rpb4-mediated coordination, showing an exceptionally prominent role for Xrn1. Consequently, we next focused our study on characterizing the specific interplay between Xrn1 and Rpb4.

**Figure 3.**
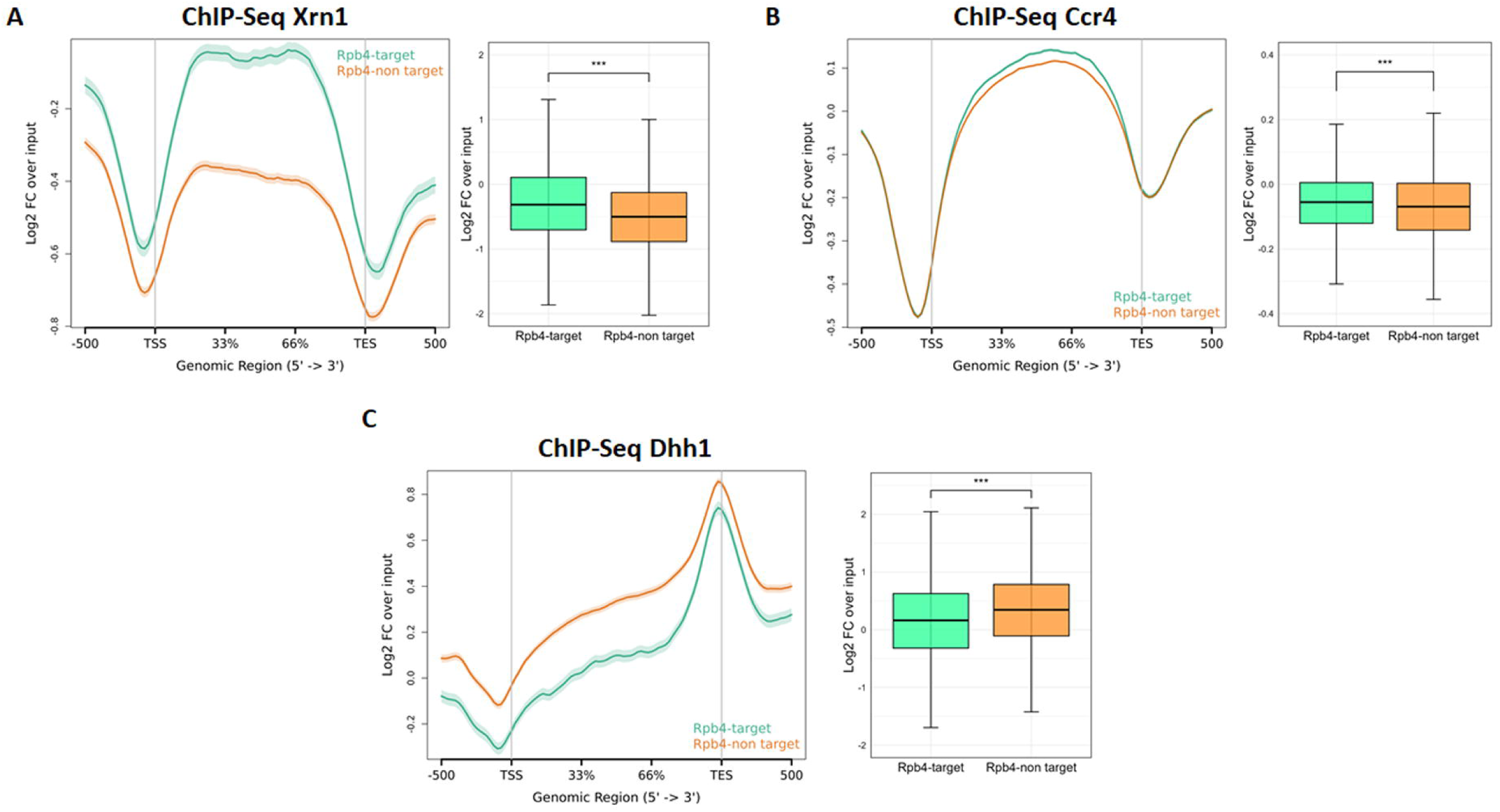
ChIP-Seq analysis of crosstalk elements occupancy. (A) Input-normalized average metagene profile of Xrn1-TAP occupancy over gene bodies and flanking regions of all protein-coding genes, categorized into Rpb4-target and non-target genes. (B) Analysis as in (A) using a Ccr4-TAP strain. (C) Analysis as in (A) using a Dhh1-TAP strain. Data represent the mean of two biological replicates. TSS, transcription start site; TES, transcription end site. Asterisks indicate statistically significant differences (* P < 0.05, ** P < 0.01, and *** P < 0.001; Wilcoxon rank-sum test). TSS, transcription start site; TES, transcription end site.

Since Rpb4-target genes exhibit higher levels of both Xrn1 and RNA pol II, we next analysed the Xrn1/Rpb3 ratio (Figure 4A). This allowed us to determine whether Xrn1 accumulation simply correlates proportionally with polymerase levels or represents a specific enrichment. As we can observe in the Figure 4A, an increase in the association of Xrn1 relative to RNA pol II for the Rpb4-target genes with respect the rest of the genes was observed, although the shape of the profile did not vary between them, suggesting that these genes are specific genomic targets of Xrn1. Notably, the ChIP-Seq Xrn1/BioGRO-Seq ratio was significantly elevated along gene body for the Rpb4-target genes (Figure 4B). This enrichment suggests that Xrn1 recruitment to these loci is not merely a passive consequence of high TR, but rather a specific targeted mechanism operating at Rpb4-regulated genes.

**Figure 4.**
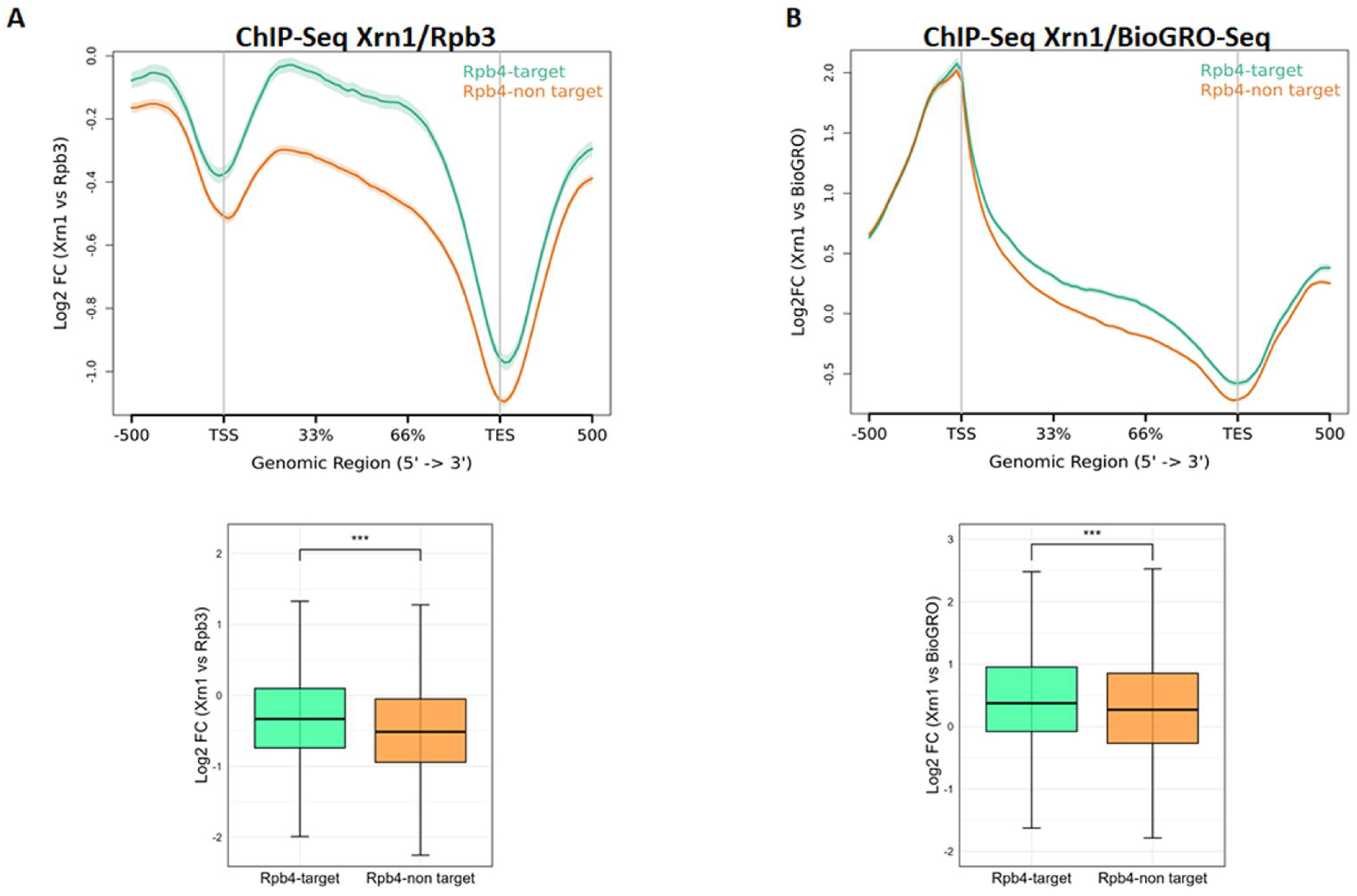
Xrn1 recruitment relative to RNA polymerase II transcription. **(A)** Metagene profile of the Xrn1/Rpb3 ratio, expressed as the log_2_ fold change of Xrn1-TAP ChIP over Rpb3 ChIP signal across gene bodies and flanking regions in Rpb4-target and non-target genes. **(B)** Average metagene profile showing the log_2_ fold change of Xrn1 occupancy over BioGRO-Seq signal across gene bodies and flanking regions in Rpb4-target and non-target genes. Values represent the average of two independent biological replicates. TSS, transcription start site; TES, transcription end site. Asterisks indicate statistically significant differences (* P < 0.05, ** P < 0.01, and *** P < 0.001; Wilcoxon rank-sum test). TSS, transcription start site; TES, transcription end site.

### The absence of Xrn1 alters Rpb3 genomic occupancy with a stronger impact on Rpb4-dependent genes

Given that genes whose mRNAs show imprinting by Rpb4 appear to be specific targets of Xrn1, we next decided to investigate the impact of Xrn1 on transcription. Specifically, we analysed RNA pol II genomic occupancy in an *xrn1*Δ mutant strain, with respect to an isogenic wild-type strain, by ChIP-Seq. As observed in Figure 5A, an *xrn1*Δ mutant exhibited a reduced RNA pol II recruitment within the 5’ end of the gene, as previously described (Begley et al., 2021; Fischer et al., 2020; Haimovich et al., 2013; Medina et al., 2014) but an increase at the 3’ end of the gene. Interestingly, the lack of Xrn1 abolishes the difference in the RNA pol II profile seen in WT cells (see also Fig. 1) suggesting that Xrn1 is necessary to maintain the distinct transcriptional regulation of Rpb4-target genes, highlighting a functional interdependence between both factors at these genes.

**Figure 5.**
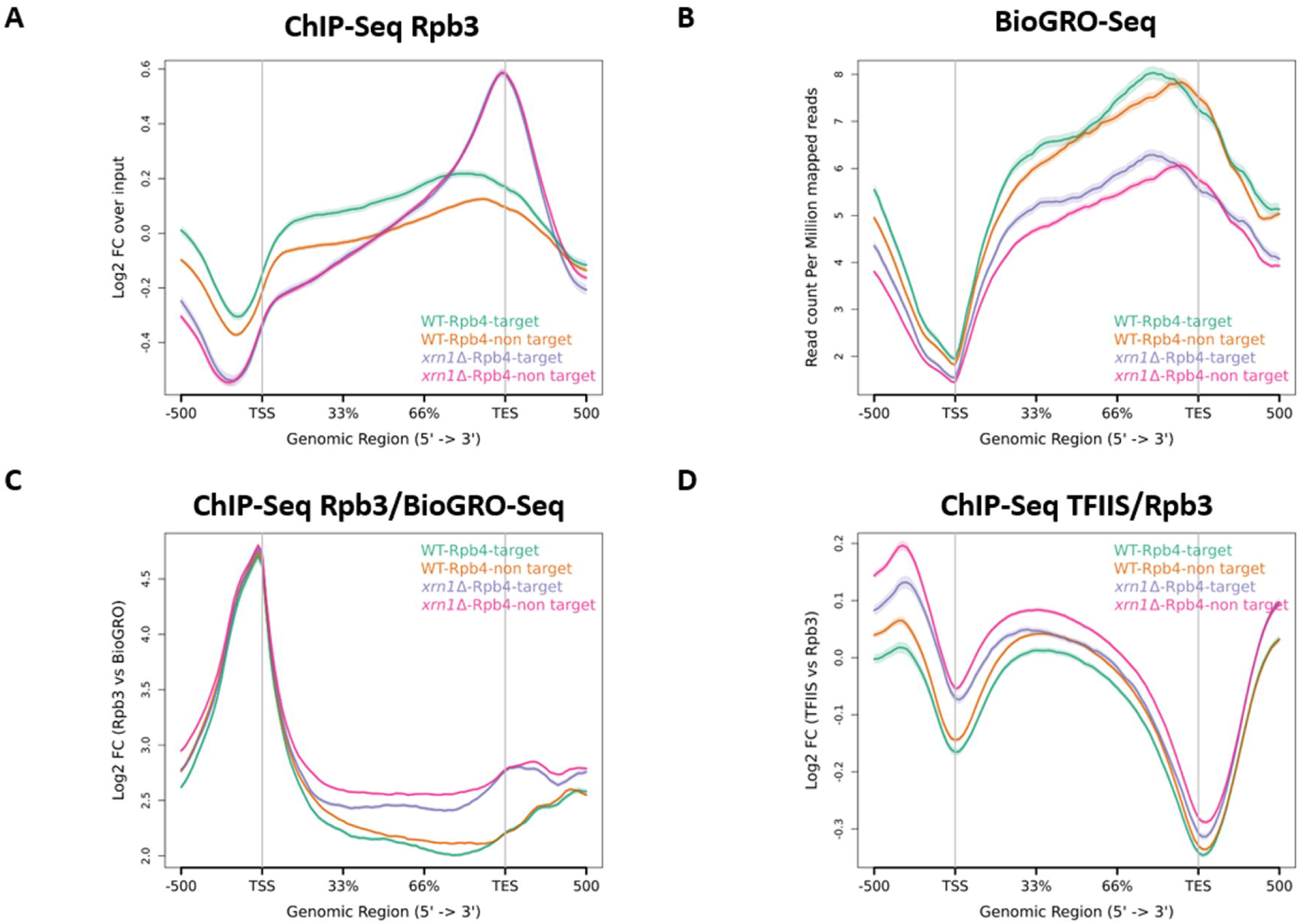
Genome-wide analysis of RNA polymerase II occupancy and transcription in the *xrn1*Δ mutant relative to wild type. (**A**) Input-normalized average metagene profile of Rpb3 over gene bodies and flanking regions of protein-coding genes, categorized into Rpb4-target and non-target genes in wild-type and *xrn1*Δ strains. **(B)** Metagene profile of BioGRO-Seq data over gene bodies and flanking regions for Rpb4-target and non-target genes in wild-type and *xrn1*Δ strains. **(C)** Average metagene profile showing the log_2_ fold change of Rpb3 occupancy over BioGRO-Seq signal across gene bodies and flanking regions in both strains. **(D)** Metagene profile of the TFIIS/Rpb3 ratio (log_2_ fold change of TFIIS ChIP over Rpb3 ChIP) across Rpb4-target and non-target genes in wild-type and *xrn1*Δ strains. WT data were previously shown in Figures 1 and 2 and are included here for reference. Results are calculated as the mean from two independent biological replicates. TSS, transcription start site; TES, transcription end site.

Next, we used previously published BioGRO-Seq data from an *xrn1*Δ mutant (Begley et al., 2021) and re-analysed the transcriptional profile of Rpb4-target and non-target genes compared with a wild-type strain. As observed in Figure 5B, the *xrn1*Δ mutant showed a global reduction in TR, as previously described (Begley et al., 2021). However, although global TR decreased, it remained significantly higher at Rpb4-target genes, mirroring the pattern observed for the wild-type strain. Moreover, the shape of the profiles is very similar to that of WT ones. These data suggest that Rpb4-target genes possess a higher TR than non-target genes, independently of the presence of Xrn1. Consistent with the literature indicating that Xrn1 deficiency provokes RNA pol II backtracking near polyadenylation sites (PAS) (Begley et al., 2021; Fischer et al., 2020), we observed a global increase in the Rpb3/BioGRO-Seq ratio upstream of the transcription termination site in the *xrn1*Δ strain, probably corresponding with PAS (Figure 5C). Interestingly, the Rpb3/BioGRO-Seq ratio remained lower at Rpb4-target genes than at non-targets even in the absence of Xrn1, suggesting that this specific subset of genes exhibits a lower baseline of polymerase backtracking independent from Xrn1. In agreement with these findings, the lack of Xrn1 led to a general increase in TFIIS recruitment along gene bodies (Figure 5D), as previously described (Begley et al., 2019). Nevertheless, Rpb4-target genes exhibited a lower TFIIS/Rpb3 ratio than the rest of the genes even in an *xrn1*Δ mutant. This confirms that Rpb4-target genes retain a reduced backtracking phenotype, even under conditions of markedly elevated global RNA pol II stalling caused by the absence of Xrn1.

### Simultaneous deletion of RPB4 and XRN1 severely impairs cell growth

Given our data indicating a functional relationship between Rpb4 and Xrn1, we next analysed the growth phenotype of the double *xrn1*Δ *rpb4*Δ mutant. For this purpose, we performed drop dilution assays at different temperatures using both single and double mutants (Figure 6). As expected, a strong genetic interaction was observed, confirming the functional interplay between both proteins. These findings point to Rpb4 and Xrn1 operating in converging pathways essential for normal cell growth under both optimal and stress conditions.

**Figure 6.**
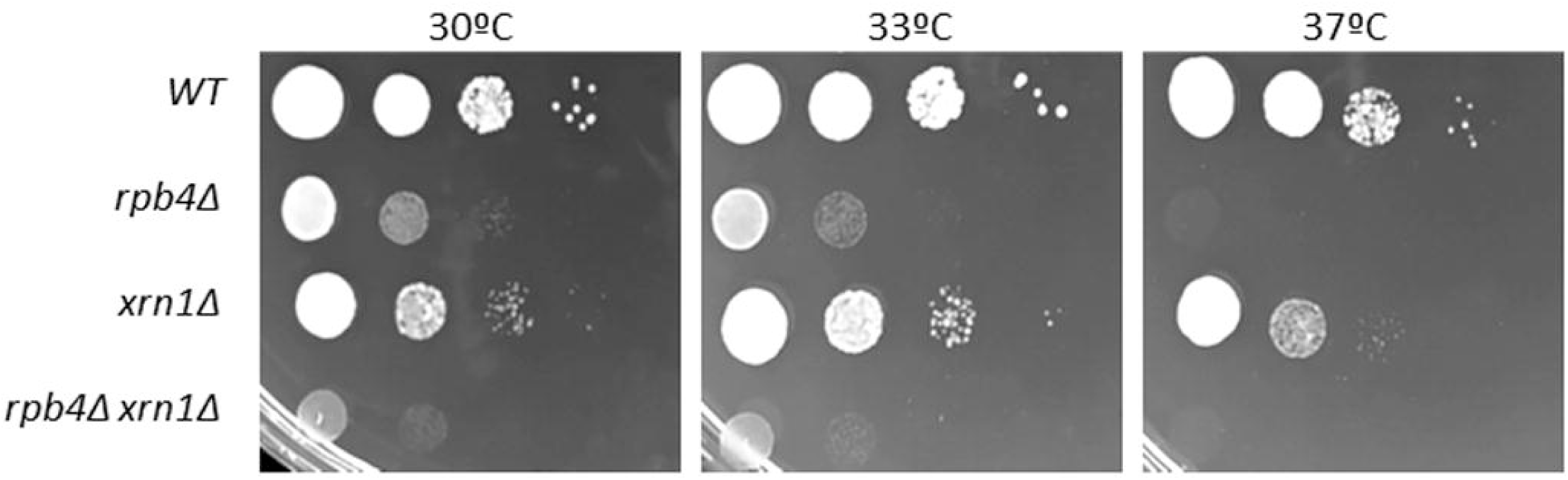
Genetic interactions between *RPB4* and *XRN1*. **S**pot assay (10-fold dilutions) of *rpb4*Δ and *xrn1*Δ single and double mutants grown at different temperatures in YPD medium. Note that the small (*xrn1*) and medium (*rpb4*) growth deficiency is strongly aggravated in a double *xnr1 rpb4* mutant.

### Loss of Rpb4 alters Xrn1 protein integrity, yet the association of Rpb4 and Xrn1 with chromatin and RNA is mutually independent

Given that Rpb4-target genes seems to be genomic targets of Xrn1, we wondered if Xrn1 absence could affect Rpb4 association to chromatin and *vice versa*. To test this, we isolated chromatin-enriched fractions using the yChEFs procedure (Cuevas-Bermúdez et al., 2019, 2020) in wild-type as well as in *xrn1*Δ and *rpb4*Δ mutant strains, and analysed by western blot the association of both Rpb4 and Xrn1 (by using specific antibodies). A specific antibody against histone H3 was used to confirm the presence of chromatin, and an anti-phosphoglycerate kinase 1 (Pgk1) antibody to rule out cytoplasmic contamination in the chromatin fractions. As shown in Figure 7A (left panel), deletion of *RPB4* did not alter the levels of Xrn1 associated with chromatin, but rather slightly increases them, as indicated by the Xrn1/H3 ratio depicted in the graph (Figure 7A, right panel). Curiously, in the absence of Rpb4, Xrn1 displayed a double-migrating band present in chromatin fractions as well as in whole-cell extracts and free-chromatin fractions. This double band might reflect a proteolytic cleavage event, suggesting that the Xrn1 protein could be more unstable in the absence of Rpb4. Additionally, Rpb4 association with chromatin was not significantly altered in the absence of Xrn1 (Figure 7A). Taken together, these data suggest that the association of Rpb4 and Xrn1 with chromatin is mutually independent, despite the functional association between both proteins.

**Figure 7.**
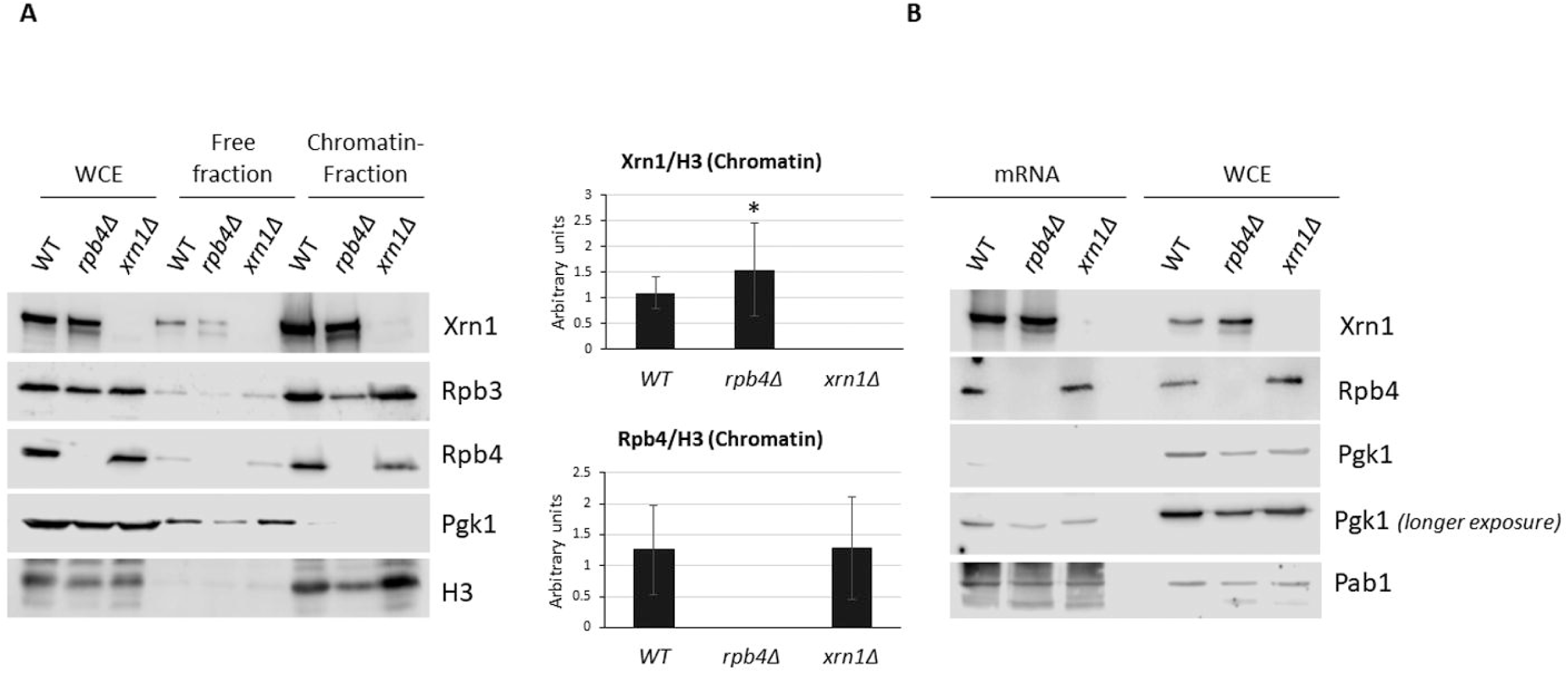
Association of Xrn1 and Rpb4 with chromatin and mRNA in *rpb4*Δ and *xrn1*Δ mutants. A) *Left*: Whole-cell extracts, chromatin-free fractions, and chromatin-associated proteins isolated using the yChEFs protocol (Cuevas-Bermúdez et al., 2019, 2020) from wild-type, *rpb4*Δ and *xrn1*Δ strains grown in YPD medium at 30 °C were analysed by Western blot with specific antibodies against Xrn1, Rpb3 and Rpb4. Histone H3 was used as a marker for chromatin-associated proteins, and Pgk1 as a control for cytoplasmic contamination. *Right:* Quantification of chromatin fractions from the left panel, showing the signal of each protein normalized to histone H3 in *rpb4*Δ and *xrn1*Δ mutants relative to the wild-type strain. Data represent the median and standard deviation of at least three independent biological replicates (P < 0.05, Student’s t-test). B) Western blot of Rpb4 and Xrn1 in whole-cell extracts and oligo(dT)-purified mRNA fractions following UV irradiation (1200 mJ/cm2 of 254 nm) in wild-type, *rpb4*Δ and *xrn1*Δ strains. Anti-Pgk1 antibody served as a negative control and anti-Pab1 antibody as a positive control for mRNA-bound proteins.

Based on the role of both proteins not only in transcription regulation but also in mRNA fate determination (Begley et al., 2019; Blasco-Moreno et al., 2019; Fischer et al., 2020; Garrido-Godino et al., 2016, 2021, 2022; Lotan et al., 2005; Medina et al., 2014), we next sought to study the mRNA association of both proteins in the *xrn1*Δ and *rpb4*Δ mutants. For this purpose, we grew cells at a permissive temperature and performed UV-light crosslinking to covalently bind the proteins to mRNA (Garrido-Godino et al., 2016). Cells were then lysed, and mRNA was isolated using magnetic oligo-dT beads prior to western blot analysis to detect mRNA-bound proteins. We used Pgk1 antibodies to detect cytoplasmic contaminant proteins, and a Pab1-specific antibody to confirm mRNA purification. As shown in Figure 7B, deletion of *RPB4* did not alter the association of Xrn1 with mRNA; likewise, deletion of *XRN1* seemed not to change the association of Rpb4 with mature polyA-mRNA either. However, because this approach specifically detects fully polyadenylated mRNA, we cannot rule out the possibility that the co-transcriptional association of these proteins with nascent mRNA might be affected.

### Rpb4 physically interacts with Xrn1 in an RNA-dependent manner

Having demonstrated that Rpb4-imprinted genes are genomic targets of Xrn1, that *XRN1* and *RPB4* genetically interact, and that Xrn1 protein integrity is altered in the absence of Rpb4, we next investigated whether these two proteins could physically contact each other. To this end, we isolated chromatin-enriched and free-chromatin fractions using the yChEFs procedure (Cuevas-Bermúdez et al., 2019, 2020), and performed protein immunoprecipitation assays using specific antibodies. Rpb4 subunit of RNA pol II, as well as the Rpb3 subunit (as an indicator of total RNA pol II) were purified, from both the free-chromatin fraction and from chromatin fractions. As shown in Figure 8, Rpb4 co-immunoprecipitated with Xrn1 in both fractions. Interestingly, this association was not observed when total RNA pol II (Rpb3) was purified, which indicate that Rpb4 and Xrn1 seem not interact via a direct association between Xrn1 and the RNA pol II complex. However, a direct protein-protein interaction between Xrn1 and complete RNA pol II could not be fully ruled out, as it has been previously described by other authors (Harlen & Churchman, 2017). Note that this methodology only focuses on a single locus under the induction conditions of the *GAL1* promoter (Harlen & Churchman, 2017). To definitively determine whether the Xrn1-Rpb4 interaction could be mediated through RNA, we performed an RNase treatment prior to the yChEFs fractionation in the same samples. Notably, under these RNA-depleted conditions, the association between Rpb4 and Xrn1 was drastically reduced.

**Figure 8.**
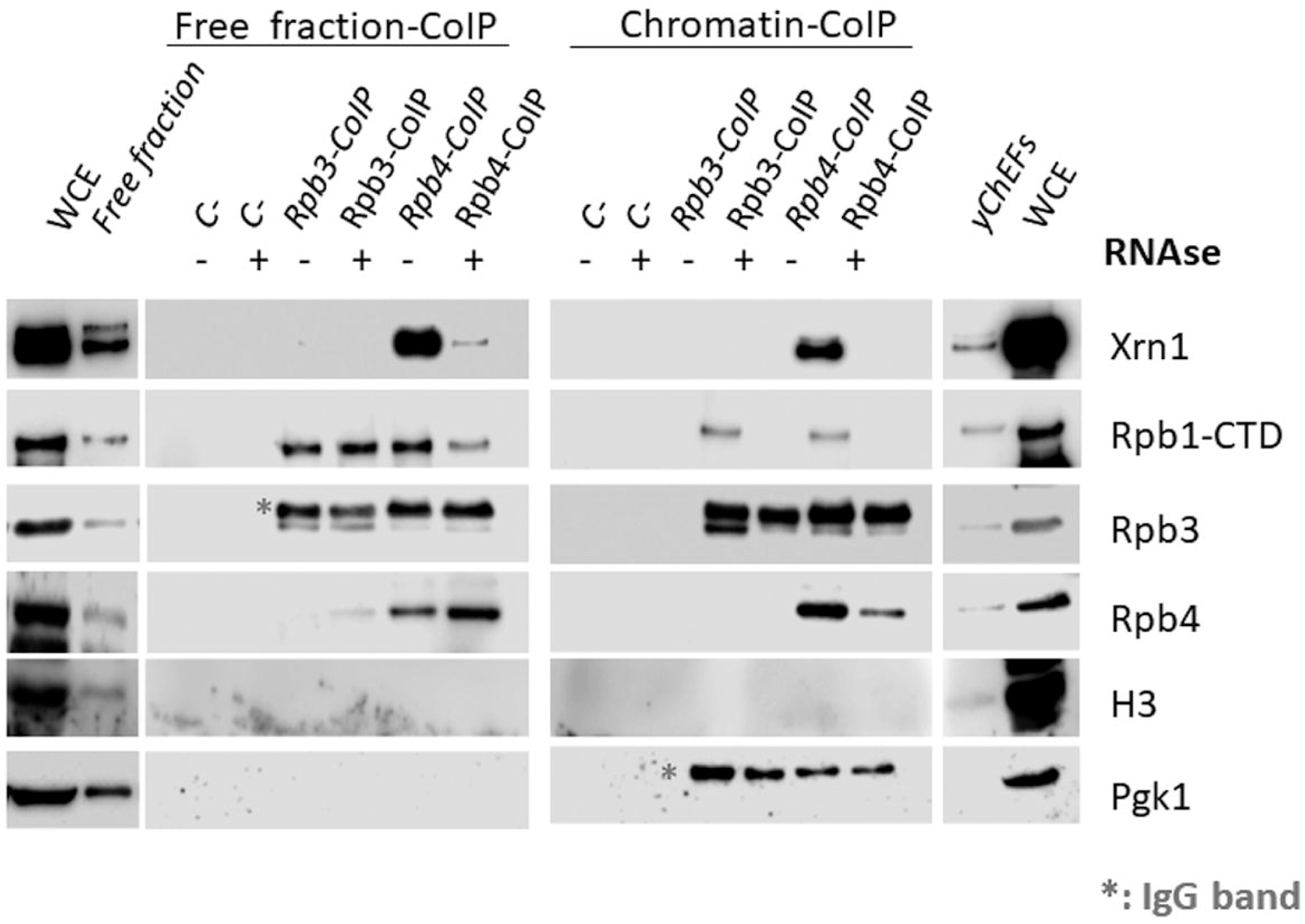
Co-immunoprecipitation of Xrn1 and Rpb4. RNA polymerase II was immunoprecipitated using anti-Rpb3 and anti-Rpb4 antibodies from chromatin-free and chromatin-enriched fractions obtained via the yChEFs protocol (Cuevas-Bermúdez et al., 2019, 2020), in a wild-type strain grown in YPD at 30 °C. Samples from both chromatin-free and chromatin-enriched fractions were analysed with and without RNase treatment. Western blot analysis was performed for Xrn1 and RNA polymerase II subunits Rpb1 (anti-CTD antibody), Rpb3, and Rpb4. Histone H3 was used as a positive control for chromatin-enriched fractions, and Pgk1 as a control for the free fraction.

Taken together, these results point to that the physical interaction between Xrn1 and Rpb4 occurs post-RNA imprinting and is mediated by RNA.

## DISCUSSION

The study is based on the hypothesis that the imprinting carried out by the RNA pol II subunits Rpb4/7 on a large set of yeast mRNAs implies a “coordinating” role for these proteins (Calvo, 2020; Garrido-Godino et al., 2022; Goler-Baron et al., 2008; Harel-Sharvit et al., 2010) and, therefore, they must interact, in some way, with other proteins that imprint different groups of genes, such as Xrn1, as we had previously demonstrated was the case with Puf3 (Garrido-Godino et al., 2021). The potential coordinating role of Rpb4/7 in mRNA imprinting could be exerted through direct or indirect mechanisms involving physical or genetic interactions, or both. In fact, the co-transcriptional imprinting of Xrn1, Rpb4 and some other dozens of proteins on yeast mRNAs has been demonstrated (Harlen & Churchman, 2017; Urim et al., 2026). However, in the study of (Urim et al., 2026), the absence of Rpb4 did not significantly affect Xrn1 imprinting, unlike what occurs with other imprinting proteins. Our study, therefore, wish to uncover the potential interdependence of Rpb4 and Xrn1 imprinting in yeast.

To test the hypothesis, we set out to determine whether Rpb4 and Xrn1 interact physically and/or genetically. We have demonstrated that Rpb4 and Xrn1 do indeed interact genetically (Figure 6) and physically in a RNA-dependent manner (Figure 8). The physical interaction, however, does not imply that the binding of one determines the binding of the other, either for mRNA or for chromatin (Fig. 7). This latest finding is therefore consistent with the aforementioned ability of Xrn1 to bind to mRNA co-transcriptionally in the absence of Rpb4, as observed by M. Choder’s group (Urim et al., 2026). The conclusion that emerges from all these results is that, although in a wild-type cell Xrn1 appears to imprint mRNAs that come into contact with Rpb4 in a co-transcriptional manner, the presence of this protein is not essential for Xrn1 to bind to mRNAs. However, we cannot exclude that absence of Rpb4 could influence functions associated to mRNA imprinting. The worsening of the growth phenotype of an *rpb4* mutant due to the absence of Xrn1 suggests that at least one of the two proteins must imprint the mRNAs to enable vital functions under optimal environmental conditions.

Regarding the model of cooperation between these two factors, we have found the following: the target genes of Xrn1 are mostly target genes of Rpb4. Xrn1 acts on its target genes by activating transcription through its roles in initiation and elongation (Begley et al., 2019, 2021; Haimovich et al., 2013). Its most evident effect is to reduce the rate of non-elongating RNA polymerase (backtracking, Fig. 5C), thereby facilitating the action of TFIIS (Fig. 5D). Rpb4 target genes also exhibit a lower backtracking rate than the rest (Fig. 2 and Fig. 5C).

Thus, it appears that both factors act in a similar manner: the presence of Rpb4 (at its targets) reduces RNA polymerase II backtracking at those genes (Fig. 5C). Similarly, the presence of Xrn1 (in the wild-type strain) also reduces backtracking because its absence increases it (Fig. 5C). In both cases, increased backtracking is associated with a higher presence of TFIIS (Fig. 5D). Xrn1, however, does not act in the same way as other elongation factors involved in imprinting, such as Dhh1 and Ccr4, likely because it preferentially binds to Rpb4 target genes (Fig. 3) in a specific manner (Fig. 4A–B). In fact, other genomic imprinting factors appear to have no effect on RNA pol II backtracking—such as Ccr4 (Begley et al., 2019)—or even to increase it in a specific group of growth-related genes, such as Sfp1 (Kelbert et al., 2024). Therefore, it appears that mRNA imprinting can occur through different molecular mechanisms.

Therefore, it would be interesting to uncover the molecular mechanism underlying the cooperation between Rpb4 and Xrn1 and what distinguishes it from other co-imprinting factors with Rpb4/7, such as Puf3 (Garrido-Godino et al., 2021) and Sfp1 (Kelbert et al., 2024), which act on a limited number of targets, Ccr4 (Begley et al., 2019; Gupta et al., 2016), which shows no preference for Rpb4 targets (this study), or the FACT complex proteins (Spt16 and Pob3), whose binding to mRNA is strictly dependent on the presence of Rpb4 (Urim et al., 2026). However, it is interesting to note that many of the mRNA imprinting factors described here (Sfp1, Rpb4, Xrn1, Ccr4, FACT) are involved in transcriptional elongation, which reinforces the idea that mRNA imprinting is primarily a co-transcriptional phenomenon in which the protein that binds to the nascent mRNA originates from the transcriptional elongation complex and dissociates from it when changes occur in the complex’s structure or activity, such as backtracking (Begley et al., 2019; Kelbert et al., 2024; Pérez-Ortín et al., 2026).

## MATERIALS AND METHODS

### Yeast strains, growth conditions, genetic manipulations and genetic analysis

Common yeast media, growth conditions and genetic techniques were used as described elsewhere (Garcia-Lopez et al., 2010). Briefly, yeast strains were grown at 30°C in liquid YPD media containing 2% glucose, unless otherwise specified. When necessary, synthetic SD media was used, supplementing it with the specific nutrients required for auxotrophic deficiencies.

YFN895 strain was obtained by chromosomal integration of a PCR product amplified using genomic DNA from strain Y04540 (Euroscarf) containing the *xrn1D::KanMX4* construction as template, in YFN679. Oligonucleotides used for amplification are detailed in Supplementary Table S2.

Yeast strains are listed in Supplementary Tables S1.

### Chromatin immunoprecipitation and sequencing analyses

For chromatin immunoprecipitation followed by high-throughput sequencing (ChIP-Seq) analysis, biological duplicates of wild-type, TAP-tagged and deletion strains were prepared, according to the protocols previously described (Begley et al., 2019; Martínez-Fernández et al., 2020). Briefly, 200 mL of yeast cultures (OD600 approx. 0.6) were fixed with 1% formaldehyde for 15 minutes, followed by quenching with 2.5 M glycine for 5 minutes. After four washes in Tris-buffered saline (150 mM NaCl, 20 mM Tris-HCl, pH 7.5), cells were resuspended in lysis buffer and disrupted using glass beads. The extracts were adjusted to 1.5 mL and sheared using a Diagenode Bioruptor (45 min, 30s cycles). Immunoprecipitation was carried out using 300 μL of the sonicated material with either Dynabeads Pan Mouse IgG (for TAP strains) or anti-Rpb3 (1Y26, Abcam) coupled to anti-mouse IgG Dynabeads. Following standard washing and decrosslinking, DNA was recovered using the Nucleospin Extract II kit (Macherey-Nagel), concentrated via lyophilisation, and sequenced on an Illumina HiSeq 2000 platform.

Quality control of the FASTQ sequencing datasets was conducted via FastQC v0.12.0 (Andrews, S., 2010), with metrics subjected to visual examination. Following the quality check, low-quality bases were trimmed from the ends of each read with Sickle v1.33 (Joshi & Fass, 2011), and any reads that consequently became too short were discarded. The sequencing adapters identified during the quality control were removed from the 3’ end of the reads, after which duplicated reads are collapsed into groups. Both processes were assessed using Fastx Clipper and Fastx Collapser from FASTX-Toolkit v0.0.14 (Gordon, A., & Hannon, G.J., 2010). Lastly, the resulting reads were mapped to she *S. cerevisiae* S288C reference genome (assembly R64; GenBank assembly accession GCF_000146045.2) with Bowtie2 v2.5.4 (Langmead & Salzberg, 2012). SAM alignment files were transformed into BAM, sorted and indexed using SamTools View, SamTools Sort and SamTools Index, respectively, from SamTools v1.23.1 (Danecek et al., 2021), and finally visually inspected with the Integrative Genomics Viewer (IGV) (Robinson et al., 2011; Thorvaldsdóttir et al., 2013). Due to the similarity of the average occupancy profiles of the two biological replicates, the BAM files from each sample were merged and indexed using SamTools Merge and SamTools Index, respectively. All bioinformatics packages were installed via Bioconda (Grüning et al., 2018).

The BioGRO-Seq samples were processed using the same pipeline described above, with the exception of the collapsing of duplicate reads, which was omitted due to the genomic nature of the sequencing.

All high-throughput sequencing datasets generated and analysed in this study are available in the NCBI Gene Expression Omnibus (GEO) database (https://www.ncbi.nlm.nih.gov/geo/). The data generated in this work (Rpb3 ChIP-Seq in WT and *xrn1*Δ strains, as well as Xrn1-TAP, Ccr4-TAP, and Dhh1-TAP ChIP-Seq) are deposited under accession number GSE354055. Previously published datasets were retrieved from GEO under entries GSE127179 (TFIIS and Rpb3 ChIPin WT and *xrn1*Δ TFIIS-HA strains) (Begley et al., 2019) and GSE153036 (BioGRO-Seq data) (Begley et al., 2021).

### Isolation of the mRNA-associated proteins

mRNA crosslinking was carried out as described in (Garrido-Godino et al., 2016) with some modifications. Briefly, 250 ml of yeast cultures (OD_600_ ∼ 0.7-0.8) were irradiated with 1200 mJ/cm^2^ of 254 nm UV light using a UV crosslinker (Biolink Shortwave 254 nm). Cells were subsequently resuspended in 1000 µl of lysis buffer (20 mM Tris pH 7.5, 0.5 M NaCl, 1 mM EDTA, 1x protease inhibitor cocktail [Complete; Roche], 0.5 mM phenylmethylsulfonyl fluoride (PMSF)) and disrupted with 500 µl of glass beads (425-600 µm, Sigma), by vortexing for 15 min at 4°C. The supernatant was then sheared using a 1-2 ml syringe and clarified by high-speed centrifugation to obtain the crude extract, from which an input aliquot was reserved. Poly(A)+ RNA and its associated proteins were captured using oligo(dT) magnetic beads (New England BioLabs, cat no. S1419S) for 30 min at room temperature. After rigorous washing to remove non-specific binders with loading buffer (20 mM Tris–HCl pH 7.5, 0.5M NaCl, 1mM EDTA, 1x protease inhibitor cocktail [Complete; Roche], 0.5 mM PMSF) and once with low salt buffer (10 mM Tris–HCl pH 7.5, 0.1 M NaCl, 1 mM EDTA), the protein-RNA complexes were eluted in 25 µl of Laemmli buffer by boiling. The eluates were resolved by SDS-PAGE and identified via Western blot using different antibodies.

### Yeast chromatin-enriched fractions preparation

Chromatin-bound proteins were extracted from 50 ml of YPD cultures (OD_600_ ∼0.6–0.8) following the yChEFs protocol (Cuevas-Bermúdez et al., 2019, 2020). Analysis of the enriched fractions was performed through Western blotting, employing the specific antibodies indicated above.

### Modified yeast chromatin-enriched fractions preparation (myChEFs)

To perform co-immunoprecipitation of chromatin-bound proteins, a modification of the yChEFs protocol termed myChEFs (modified yChEFs) was used. Briefly, 200 ml of *Saccharomyces cerevisiae* cultures were grown to exponential phase (OD_600_ ∼0.7–0.8). The standard yChEFs purification was performed to obtain the chromatin-enriched pellet (Cuevas-Bermúdez et al., 2019, 2020). The resulting pellet was subsequently resuspended in 10 mM Tris-HCl pH 7.5, 2.5 mM MgCl_2_, 0.1 mM CaCl_2_) supplemented with 150 U DNAse I (18047019 Invitrogen) for enzymatic digestion. The suspension was incubated for 2 h at 4°C with constant rotation on a wheel.

Whenever required RNase (2 mg/ml) was added to WCE prior chromatin isolation, and was incubated for 1 h at 37 °C as previously described (Cuevas-Bermúdez et al., 2019).

The resulting material was then used for protein immunoprecipitation using anti-Rpb3 antibody (anti-POLR2C;1Y26, Abcam) or anti-Rpb4 (Pol II RPB4 (2Y14); Biolegend) coupled to 100 µl of Dynabeads Sheep-anti-Mouse IgG (Invitrogen) per sample. Protein immunoprecipitation protocol was performed as previously detail (Garrido-Godino et al., 2013). The affinity-purified proteins were released from the beads by boiling for 10 min and were analysed by western blotting with different antibodies.

### SDS-PAGE and Western blot analysis

Protein electrophoresis and western blot were carried out as described in (Garrido-Godino et al., 2013).

For the western blot analyses the following antibodies were used: anti-CTD (Cuevas-Bermúdez et al., 2019), anti-Rpb3 (anti-POLR2C;1Y26, Abcam), anti-Rpb4 (Pol II RPB4 (2Y14); Biolegend), anti-phosphoglycerate kinase, Pgk1 (22C5D8; Invitrogen), anti-H3 (ab1791; Abcam), anti-Xrn1 (gift from Arlen Johnson), anti-Ccr4 (gift from Martine Collart) and anti-Pab1 (MCA-1G1, EnCor Biotechnology).

## Bibliography

1. Andrews, S. (2010). FastQC A Quality Control tool for High Throughput Sequence Data. https://www.bioinformatics.babraham.ac.uk/projects/fastqc/

2. Babbarwal, V., Fu, J., & Reese, J. C. (2014). The Rpb4/7 module of RNA polymerase II is required for carbon catabolite repressor protein 4-negative on TATA (Ccr4-not) complex to promote elongation. Journal of Biological Chemistry, 289(48), 33125–33130. 10.1074/jbc.C114.601088

3. Begley, V., Corzo, D., Jordán-Pla, A., Cuevas-Bermúdez, A., Miguel-Jiménez, L. de, Pérez-Aguado, D., Machuca-Ostos, M., Navarro, F., Chávez, M. J., & Pérez-Ortín, J. E. (2019). The mRNA degradation factor Xrn1 regulates transcription elongation in parallel to Ccr4. Nucleic acids research, 47(18), 9524–9541. 10.1093/nar/gkz660

4. Begley, V., Jordán-Pla, A., Peñate, X., Garrido-Godino, A. I., Challal, D., Cuevas-Bermúdez, A., Mitjavila, A., Barucco, M., Gutiérrez, G., & Singh, A. (2021). Xrn1 influence on gene transcription results from the combination of general effects on elongating RNA pol II and gene-specific chromatin configuration. RNA biology, 18(9), 1310–1323. 10.1080/15476286.2020.1845504

5. Blasco-Moreno, B., de Campos-Mata, L., Böttcher, R., García-Martínez, J., Jungfleisch, J., Nedialkova, D. D., Chattopadhyay, S., Gas, M.-E., Oliva, B., & Pérez-Ortín, J. E. (2019). The exonuclease Xrn1 activates transcription and translation of mRNAs encoding membrane proteins. Nature communications, 10(1), 1298. 10.1038/s41467-019-09199-6

6. Bloom, A. L. M., Solomons, J. T. G., Havel, V. E., & Panepinto, J. C. (2013). Uncoupling of mRNA synthesis and degradation impairs adaptation to host temperature in C ryptococcus neoformans. Molecular microbiology, 89(1), 65–83. 10.1111/mmi.12258

7. Bryll, A. R., & Peterson, C. L. (2023). The circular logic of mRNA homeostasis. Transcription, 14(1-2), 18–26. 10.1080/21541264.2023.2183684

8. Calvo, O. (2020). RNA polymerase II phosphorylation and gene looping: New roles for the Rpb4/7 heterodimer in regulating gene expression. Curr Genet, 66(5), 927–937. 10.1007/s00294-020-01084-w

9. Chattopadhyay, S., Garcia-Martinez, J., Haimovich, G., Fischer, J., Khwaja, A., Barkai, O., Chuartzman, S. G., Schuldiner, M., Elran, R., Rosenberg, M. I., Urim, S., Deshmukh, S., Bohnsack, K. E., Bohnsack, M. T., Perez-Ortin, J. E., & Choder, M. (2022). RNA-controlled nucleocytoplasmic shuttling of mRNA decay factors regulates mRNA synthesis and a novel mRNA decay pathway. Nature Communications, 13(1), 7184. 10.1038/s41467-022-34417-z

10. Chávez, S., García-Martínez, J., Delgado-Ramos, L., & Pérez-Ortín, J. E. (2016). The importance of controlling mRNA turnover during cell proliferation. Current Genetics, 62(4), 701–710. 10.1007/s00294-016-0594-2

11. Chernyakov, I., Whipple, J. M., Kotelawala, L., Grayhack, E. J., & Phizicky, E. M. (2008). Degradation of several hypomodified mature tRNA species in *Saccharomyces cerevisiae* is mediated by Met22 and the 5′–3′ exonucleases Rat1 and Xrn1. Genes & Development, 22(10), 1369–1380. 10.1101/gad.1654308

12. Cheung, A. C., & Cramer, P. (2011). Structural basis of RNA polymerase II backtracking, arrest and reactivation. Nature, 471(7337), 249–253. 10.1038/nature09785

13. Choder, M. (2011). mRNA imprinting: Additional level in the regulation of gene expression. Cell Logist, 1(1), 37–40. 10.4161/cl.1.1.14465

14. Cuevas-Bermúdez, A., Garrido-Godino, A. I., Gutiérrez-Santiago, F., & Navarro, F. (2020). A Yeast Chromatin-enriched Fractions Purification Approach, yChEFs, from *Saccharomyces cerevisiae*. Bio-Protocol, 10(1). 10.21769/BioProtoc.3471

15. Cuevas-Bermúdez, A., Garrido-Godino, A. I., & Navarro, F. (2019). A novel yeast chromatin-enriched fractions purification approach, yChEFs, for the chromatin-associated protein analysis used for chromatin-associated and RNA-dependent chromatin-associated proteome studies from Saccharomyces cerevisiae. Gene Reports, 16, 100450. 10.1016/j.genrep.2019.100450

16. Dahan, N., & Choder, M. (2013). The eukaryotic transcriptional machinery regulates mRNA translation and decay in the cytoplasm. Biochim Biophys Acta, 1829(1), 169–173. 10.1016/j.bbagrm.2012.08.004

17. Danecek, P., Bonfield, J. K., Liddle, J., Marshall, J., Ohan, V., Pollard, M. O., Whitwham, A., Keane, T., McCarthy, S. A., Davies, R. M., & Li, H. (2021). Twelve years of SAMtools and BCFtools. GigaScience, 10(2), giab008. 10.1093/gigascience/giab008

18. Dori-Bachash, M., Shema, E., & Tirosh, I. (2011). Coupled evolution of transcription and mRNA degradation. PLoS biology, 9(7), e1001106. 10.1371/%20journal.pbio.1001106

19. Duek, L., Barkai, O., Elran, R., Adawi, I., & Choder, M. (2018). Dissociation of Rpb4 from RNA polymerase II is important for yeast functionality. PloS one, 13(10), e0206161. 10.1371/journal.pone.0206161

20. Duncan-Lewis, C., Hartenian, E., King, V., & Glaunsinger, B. A. (2021). Cytoplasmic mRNA decay represses RNA polymerase II transcription during early apoptosis. eLife, 10, e58342. 10.7554/eLife.58342

21. Edwards, A. M., Kane, C. M., Young, R. A., & Kornberg, R. D. (1991). Two dissociable subunits of yeast RNA polymerase II stimulate the initiation of transcription at a promoter in vitro. J Biol Chem, 266(1), 71–75.

22. Farago, M., Nahari, T., Hammel, C., Cole, C. N., & Choder, M. (2003). Rpb4p, a subunit of RNA polymerase II, mediates mRNA export during stress. Mol Biol Cell, 14(7), 2744–2755. 10.1091/mbc.e02-11-0740

23. Fischer, J., Song, Y. S., Yosef, N., di Iulio, J., Churchman, L. S., & Choder, M. (2020). The yeast exoribonuclease Xrn1 and associated factors modulate RNA polymerase II processivity in 5 ‘and 3 ‘gene regions. Journal of Biological Chemistry, 295(33), 11435–11454. 10.1074/jbc.RA120.013426

24. Freeberg, M. A., Han, T., Moresco, J. J., Kong, A., Yang, Y. C., Lu, Z. J., Yates, J. R., & Kim, J. K. (2013). Pervasive and dynamic protein binding sites of the mRNA transcriptome in *Saccharomyces cerevisiae*. Genome Biol, 14(2), R13. 10.1186/gb-2013-14-2-r13

25. Garcia-Lopez, M. C., Miron-Garcia, M. C., Garrido-Godino, A. I., Mingorance, C., & Navarro, F. (2010). Overexpression of SNG1 causes 6-azauracil resistance in *Saccharomyces cerevisiae*. Curr Genet, 56(3), 251–263. 10.1007/s00294-010-0297-z

26. García-Martínez, J., Pérez-Martínez, M. E., Pérez-Ortín, J. E., & Alepuz, P. (2021). Recruitment of Xrn1 to stress-induced genes allows efficient transcription by controlling RNA polymerase II backtracking. RNA Biology, 18(10), 1458–1474. 10.1080/15476286.2020.1857521

27. García-Martínez, José, Medina, Daniel A, Bellvís, Pablo, Sun, M., Cramer, P., Chávez, S., & Pérez-Ortín, J. E. (2021). The total mRNA concentration buffering system in yeast is global rather than gene-specific. RNA, 27(10), 1281–1290. 10.1261/rna.078774.121

28. Garrido-Godino, A. I., García-López, M. C., García-Martínez, J., Pelechano, V., Medina, D. A., Pérez-Ortín, J. E., & Navarro, F. (2016). Rpb1 foot mutations demonstrate a major role of Rpb4 in mRNA stability during stress situations in yeast. Biochimica et Biophysica Acta (BBA)-Gene Regulatory Mechanisms, 1859(5), 731–743. 10.1016/j.bbagrm.2016.03.008

29. Garrido-Godino, A. I., Garcia-Lopez, M. C., & Navarro, F. (2013). Correct Assembly of RNA Polymerase II Depends on the Foot Domain and Is Required for Multiple Steps of Transcription in *Saccharomyces cerevisiae*. Mol Cell Biol, 33(18), 3611–3626. 10.1128/MCB.00262-13

30. Garrido-Godino, A. I., Gupta, I., Gutiérrez-Santiago, F., Martínez-Padilla, A. B., Alekseenko, A., Steinmetz, L. M., Pérez-Ortín, J. E., Pelechano, V., & Navarro, F. (2021). Rpb4 and Puf3 imprint and post-transcriptionally control the stability of a common set of mRNAs in yeast. RNA biology, 18(8), 1206–1220. 10.1080/15476286.2020.1839229

31. Garrido-Godino, A. I., Martín-Expósito, M., Gutiérrez-Santiago, F., Perez-Fernandez, J., & Navarro, F. (2022). Rpb4/7, a key element of RNA pol II to coordinate mRNA synthesis in the nucleus with cytoplasmic functions in *Saccharomyces cerevisiae*. Biochimica et Biophysica Acta (BBA)-Gene Regulatory Mechanisms, 1865(5), 194846. 10.1016/j.bbagrm.2022.194846

32. Geerlings, T. H., Vos, J. C., & Raué, H. A. (2000). The final step in the formation of 25S rRNA in *Saccharomyces cerevisiae* is performed by 5’—>3’ exonucleases. RNA, 6(12), 1698–1703. 10.1017/s1355838200001540

33. Gerber, A. P., Herschlag, D., & Brown, P. O. (2004). Extensive association of functionally and cytotopically related mRNAs with Puf family RNA-binding proteins in yeast. PLoS biology, 2(3), e79. 10.1371/journal.pbio.0020079

34. Goler-Baron, V., Selitrennik, M., Barkai, O., Haimovich, G., Lotan, R., & Choder, M. (2008). Transcription in the nucleus and mRNA decay in the cytoplasm are coupled processes. Genes Dev, 22(15), 2022–2027. 10.1101/gad.473608

35. Gomez-Herreros, F., de Miguel-Jimenez, L., Millan-Zambrano, G., Penate, X., Delgado-Ramos, L., Munoz-Centeno, M. C., & Chavez, S. (2012). One step back before moving forward: Regulation of transcription elongation by arrest and backtracking. FEBS Lett, 586(18), 2820–2825. 10.1016/j.febslet.2012.07.030

36. Gomez-Herreros, F., de Miguel-Jimenez, L., Morillo-Huesca, M., Delgado-Ramos, L., Munoz-Centeno, M. C., & Chavez, S. (2012). TFIIS is required for the balanced expression of the genes encoding ribosomal components under transcriptional stress. Nucleic Acids Res, 40(14), 6508–6519. 10.1093/nar/gks340

37. Gordon, A., & Hannon, G.J. (2010). FASTX-Toolkit: FASTQ/A short-reads pre-processing tools. http://hannonlab.cshl.edu/fastx_toolkit/

38. Grüning, B., Dale, R., Sjödin, A., Chapman, B. A., Rowe, J., Tomkins-Tinch, C. H., Valieris, R., Köster, J., & Bioconda Team. (2018). Bioconda: Sustainable and comprehensive software distribution for the life sciences. Nature Methods, 15(7), 475–476. 10.1038/s41592-018-0046-7

39. Gupta, I., Clauder Münster, S., Klaus, B., Järvelin, A. I., Aiyar, R. S., Benes, V., Wilkening, S., Huber, W., Pelechano, V., & Steinmetz, L. M. (2014). Alternative polyadenylation diversifies post transcriptional regulation by selective RNA–protein interactions. Molecular systems biology, 10(2), 719. 10.1002/msb.135068

40. Gupta, I., Villanyi, Z., Kassem, S., Hughes, C., Panasenko, O. O., Steinmetz, L. M., & Collart, M. A. (2016). Translational capacity of a cell is determined during transcription elongation via the Ccr4-Not complex. Cell reports, 15(8), 1782–1794. 10.1016/j.celrep.2016.04.055

41. Haimovich, G., Medina, D. A., Causse, S. Z., Garber, M., Millan-Zambrano, G., Barkai, O., Chavez, S., Perez-Ortin, J. E., Darzacq, X., & Choder, M. (2013). Gene expression is circular: Factors for mRNA degradation also foster mRNA synthesis. Cell, 153(5), 1000–1011. 10.1016/j.cell.2013.05.012

42. Harel-Sharvit, L., Eldad, N., Haimovich, G., Barkai, O., Duek, L., & Choder, M. (2010). RNA Polymerase II Subunits Link Transcription and mRNA Decay to Translation. Cell, 143(4), 552–563. 10.1016/j.cell.2010.10.033

43. Harlen, K. M., & Churchman, L. S. (2017). Subgenic Pol II interactomes identify region specific transcription elongation regulators. Molecular systems biology, 13(1), 900. 10.15252/msb.20167279

44. Hartenian, E., & Glaunsinger, B. A. (2019). Feedback to the central dogma: Cytoplasmic mRNA decay and transcription are interdependent processes. Critical Reviews in Biochemistry and Molecular Biology, 54(4), 385–398. 10.1080/10409238.2019.1679083

45. Izban, M. G., & Luse, D. S. (1992). The RNA polymerase II ternary complex cleaves the nascent transcript in a 3’—5’ direction in the presence of elongation factor SII. Genes Dev, 6(7), 1342–1356. 10.1101/gad.6.7.1342

46. Jordán-Pla, A., Zhang, Y., García-Martínez, J., Chattopadhyay, S., Forte, A., Choder, M., Pelechano, V., & Pérez-Ortín, J. E. (2025). Proper 5’-3’ cotranslational mRNA decay in yeast requires import of Xrn1 to the nucleus. PloS One, 20(1), e0308195. 10.1371/journal.pone.0308195

47. Joshi, N. A., & Fass, J. N. (2011). Sickle: A sliding-window, adaptive, quality-based trimming tool for FastQ files. http://github.com/mloesch/sickle

48. Kelbert, M., Jordán-Pla, A., de Miguel-Jiménez, L., García-Martínez, J., Selitrennik, M., Guterman, A., Henig, N., Granneman, S., Pérez-Ortín, J. E., Chávez, S., & Choder, M. (2024). The zinc-finger transcription factor Sfp1 imprints specific classes of mRNAs and links their synthesis to cytoplasmic decay. eLife, 12, RP90766. 10.7554/eLife.90766

49. Kershaw, C. J., Costello, J. L., Talavera, D., Rowe, W., Castelli, L. M., Sims, P. F., Grant, C. M., Ashe, M. P., Hubbard, S. J., & Pavitt, G. D. (2015). Integrated multi-omics analyses reveal the pleiotropic nature of the control of gene expression by Puf3p. Scientific reports, 5, 15518. 10.1038/srep15518

50. Langeberg, C. J., Welch, W. R. W., McGuire, J. V., Ashby, A., Jackson, A. D., & Chapman, E. G. (2020). Biochemical Characterization of Yeast Xrn1. Biochemistry, 59(15), 1493–1507. 10.1021/acs.biochem.9b01035

51. Langmead, B., & Salzberg, S. L. (2012). Fast gapped-read alignment with Bowtie 2. Nature Methods, 9(4), 357–359. 10.1038/nmeth.1923

52. Larimer, F. W., Hsu, C. L., Maupin, M. K., & Stevens, A. (1992). Characterization of the XRN1 gene encoding a 5’-->3’ exoribonuclease: Sequence data and analysis of disparate protein and mRNA levels of gene-disrupted yeast cells. Gene, 120(1), 51–57. 10.1016/0378-1119(92)90008-d

53. Lotan, R., Bar-On, V. G., Harel-Sharvit, L., Duek, L., Melamed, D., & Choder, M. (2005). The RNA polymerase II subunit Rpb4p mediates decay of a specific class of mRNAs. Genes Dev, 19(24), 3004–3016. 10.1101/gad.353205

54. Martínez-Fernández, V., Cuevas-Bermúdez, A., Gutiérrez-Santiago, F., Garrido-Godino, A. I., Rodríguez-Galán, O., Jordán-Pla, A., Lois, S., Triviño, J. C., de la Cruz, J., & Navarro, F. (2020). Prefoldin-like Bud27 influences the transcription of ribosomal components and ribosome biogenesis in *Saccharomyces cerevisiae*. RNA, 26(10), 1360–1379. 10.1261/rna.075507.120

55. Medina, D. A., Jordán-Pla, A., Millán-Zambrano, G., Chávez, S., Choder, M., & Pérez-Ortín, J. E. (2014). Cytoplasmic 5′-3′ exonuclease Xrn1p is also a genome-wide transcription factor in yeast. Frontiers in genetics, 5(1). 10.3389/fgene.2014.00001

56. Miller, J. E., & Reese, J. C. (2012). Ccr4-Not complex: The control freak of eukaryotic cells. Crit Rev Biochem Mol Biol, 47(4), 315–333. 10.3109/10409238.2012.667214

57. Orlicky, S. M., Tran, P. T., Sayre, M. H., & Edwards, A. M. (2001). Dissociable Rpb4-Rpb7 subassembly of RNA polymerase II binds to single-strand nucleic acid and mediates a post-recruitment step in transcription initiation. J Biol Chem, 276(13), 10097–10102. 10.1074/jbc.M003165200

58. Parker, R. (2012). RNA degradation in Saccharomyces cerevisae. Genetics, 191(3), 671–702. 10.1534/genetics.111.137265

59. Parker, R., & Song, H. (2004). The enzymes and control of eukaryotic mRNA turnover. Nature Structural & Molecular Biology, 11(2), 121–127. 10.1038/nsmb724

60. Pelechano, V., Wei, W., & Steinmetz, L. M. (2015). Widespread Co-translational RNA Decay Reveals Ribosome Dynamics. Cell, 161(6), 1400–1412. 10.1016/j.cell.2015.05.008

61. Pérez-Ortín, J. E., Alepuz, P., Chávez, S., & Choder, M. (2013). Eukaryotic mRNA Decay: Methodologies, Pathways, and Links to Other Stages of Gene Expression. RNA Metabolism: Interactions Matter, 425(20), 3750–3775. 10.1016/j.jmb.2013.02.029

62. Pérez-Ortín, J. E., Chávez, S., & Choder, M. (2026). RNA imprinting by zinc-finger proteins. Trends in Biochemical Sciences, 51(8), 761–773. 10.1016/j.tibs.2026.04.012

63. Pérez-Ortín, J. E., Medina, D. A., Chávez, S., & Moreno, J. (2013). What do you mean by transcription rate? BioEssays, 35(12), 1056–1062. 10.1002/bies.201300057

64. Pérez-Ortín, J. E., Tordera, V., & Chávez, S. (2019). Homeostasis in the Central Dogma of Molecular Biology: The importance of mRNA instability. RNA biology, 16(12), 1659–1666. 10.1080/15476286.2019.1655352

65. Robinson, J. T., Thorvaldsdóttir, H., Winckler, W., Guttman, M., Lander, E. S., Getz, G., & Mesirov, J. P. (2011). Integrative genomics viewer. Nature Biotechnology, 29(1), 24–26. 10.1038/nbt.1754

66. Runner, V. M., Podolny, V., & Buratowski, S. (2008). The Rpb4 subunit of RNA polymerase II contributes to cotranscriptional recruitment of 3’ processing factors. Mol Cell Biol, 28(6), 1883–1891. 10.1128/MCB.01714-07

67. Schulz, D., Pirkl, N., Lehmann, E., & Cramer, P. (2014). Rpb4 subunit functions mainly in mRNA synthesis by RNA polymerase II. Journal of Biological Chemistry, 289(25), 17446–17452. 10.1074/jbc.M114.568014

68. Stevens, A. (2001). 5’-exoribonuclease 1: Xrn1. Methods in Enzymology, 342, 251–259. 10.1016/s0076-6879(01)42549-3

69. Thorvaldsdóttir, H., Robinson, J. T., & Mesirov, J. P. (2013). Integrative Genomics Viewer (IGV): High-performance genomics data visualization and exploration. Briefings in Bioinformatics, 14(2), 178–192. 10.1093/bib/bbs017

70. Timmers, H. T. M., & Tora, L. (2018). Transcript Buffering: A Balancing Act between mRNA Synthesis and mRNA Degradation. Molecular Cell, 72(1), 10–17. 10.1016/j.molcel.2018.08.023

71. Tucker, M., Staples, R. R., Valencia-Sanchez, M. A., Muhlrad, D., & Parker, R. (2002). Ccr4p is the catalytic subunit of a Ccr4p/Pop2p/Notp mRNA deadenylase complex in *Saccharomyces cerevisiae*. Embo J, 21(6), 1427–1436. 10.1093/emboj/21.6.1427

72. Urim, S., Artamov, A., Deshmukh, S. B., & Choder, M. (2026). mRNA Imprinting: Transcription apparatus can remotely control cytoplasmic post-transcriptional mechanisms by dozens of proteins. eLife, 15. 10.7554/eLife.110706.1

73. Van Dijk, E. L., Chen, C. L., d’Aubenton-Carafa, Y., Gourvennec, S., Kwapisz, M., Roche, V., Bertrand, C., Silvain, M., Legoix-Ne, P., & Loeillet, S. (2011). XUTs are a class of Xrn1-sensitive antisense regulatory non-coding RNA in yeast. Nature, 475(7354), 114–117. 10.1038/nature10118

74. Verma-Gaur, J., Rao, S. N., Taya, T., & Sadhale, P. (2008). Genomewide recruitment analysis of Rpb4, a subunit of polymerase II in *Saccharomyces cerevisiae*, reveals its involvement in transcription elongation. Eukaryot Cell, 7(6), 1009–1018. 10.1128/EC.00057-08

75. Villanyi, Z., Ribaud, V., Kassem, S., Panasenko, O. O., Pahi, Z., Gupta, I., Steinmetz, L., Boros, I., & Collart, M. A. (2014). The Not5 subunit of the ccr4-not complex connects transcription and translation. PLoS genetics, 10(10), e1004569. 10.1371/journal.pgen.1004569

76. Wilinski, D., Buter, N., Klocko, A. D., Lapointe, C. P., Selker, E. U., Gasch, A. P., & Wickens, M. (2017). Recurrent rewiring and emergence of RNA regulatory networks. Proceedings of the National Academy of Sciences, 114(14), E2816–E2825. 10.1073/pnas.1617777114

