## Supplemental Table 1 for "The Xrn1-Rpb4 axis controls mRNA synthesis and imprinting"

**Supplementary Table S1. *Saccharomyces cerevisiae* strains**

| **Strain** | **Genotype** | **Origin** |
| --- | --- | --- |
| BY4741 | *MATa his3-Δ1 leu2-Δ0 met15-Δ0 ura3-Δ0* | Euroscarf |
| Y04540 | *MATa his3-Δ1 leu2-Δ0 met15-Δ0 ura3-Δ0 xrn1Δ::KanMX4* | Euroscarf |
| XRN1-TAP | *MATa his3-Δ1 leu2-Δ0 met15-Δ0 ura3-Δ0 xrn1::XRN1-TAP-URA* | Lab Patrick Cramer |
| CCR4-TAP | *MATa his3-Δ1 leu2-Δ0 met15-Δ0 ura3-Δ0 CCR4-TAP::HIS3* | Open Biosystems |
| DHH1-TAP | *MATa his3-Δ1 leu2-Δ0 met15-Δ0 ura3-Δ0 DHH1-TAP::HIS3* | Open Biosystems |
| Y01285 | *MATa his3-Δ1 leu2-Δ0 met15-Δ0 ura3-Δ0 rpb4Δ::kanMX4* | Euroscarf |
| YFN679 | *MATa his3-Δ1 leu2-Δ0 met15-Δ0 ura3-Δ0 rpb4Δ::HIS3* | This work |
| YFN895 | *MATa his3-Δ1 leu2-Δ0 met15-Δ0 ura3-Δ0 rpb4Δ::HIS3 xrn1Δ::KANMX4* | This work |

**Supplementary Table S2. Oligonucleotides used**

| **DNA region** | **Sequence** |  |
| --- | --- | --- |
| *XRN1*-301 | ACCCTTTGACAATCCCCATTTG | Downstream of STOP codon |
| *XRN1*-501 | ACACTTGTAACAACAGCAGCA | Upstream of STOP codon |
